# A Phantom System Designed to Assess the Effects of Membrane Lipids on Water Proton Relaxation

**DOI:** 10.1101/387845

**Authors:** Oshrat Shtangel, Aviv A. Mezer

## Abstract

**Purpose:** Quantitative magnetic resonance imaging (qMRI) provides a method for the non-invasive study of brain structure and associated changes, expressed in physical units. qMRI parameters have been shown to reflect brain tissue composition such as myelin. Nevertheless, it remains a major challenge to identify and quantify the contributions of specific molecular components to the MRI signal. Here, we describe a phantom system that can be used to evaluate the contribution of human brain lipids to qMRI parameters.

**Methods:** A thin layer evaporation-hydration technique was used to formulate liposomes that mimic the physiological bi-layered membrane lipid environment. We then applied quantitative clinical MRI techniques with adjusted bias corrections in order to test the ability of the phantom system to estimate multiple qMRI parameters such as proton density (PD), T1, T2, T2* and magnetization transfer (MT).

**Results:** The results indicated that phantoms composed of various lipids could provide a stable and reliable estimation of qMRI parameters. In addition, the calculated water fraction (WF) maps for the phantoms were found to accurately represent the true WF volumes.

**Conclusion:** We have successfully created a biologically relevant liposome phantom system whose lipid composition can be fully controlled. This system can be used to measure the contributions of lipids to qMRI parameters under conditions that are relevant to in-vivo human scans.

## Introduction

Magnetic resonance imaging (MRI) is a fundamental technique that is often used for clinical assessment and structural research of the human brain. Quantitative MRI (qMRI) analysis produces parametric maps that have measurable units such as relaxation times (s). These qMRI maps enable a reliable comparison of brain state across different time points and MRI scanners, making it possible to assess normal brain development, as well as pathological conditions^1^.

qMRI maps also have several limitations, including image resolution, and ambiguous interpretation of the signal received. While brain cellular structures are on a micrometer scale, qMRI maps have only millimeter resolution. Furthermore, the MRI signal is driven by the hydrogen protons of water molecules, and the effects of the biological environments surrounding the protons are difficult to estimate^2^. Therefore, tools with the ability to separate the signal to its components are of extreme value.

Different qMRI parameters have been associated with a number of biological sources. For example, changes in myelination levels are reflected by alterations in T1, MT, and T2 ^3-6^. Furthermore, the iron content and water fraction (WF) of cellular compartments, are also thought to influence the qMRI parameters^7,8^. Currently, it still remains a major challenge to localize and quantify the contribution of a specific biochemical environment to an MRI signal in vivo.

In attempts to resolve this issue, great efforts in the qMRI field have been invested in building phantoms that can mimic the natural environment of the human brain. Such phantoms permit insights into potential biological sources of the signal while maintaining strict control over the precise content of the phantom^9^. Another critical advantage is that the use of phantoms can ensure that the qMRI parameters are consistent and reliable in time and space. The main objective of this project was to establish a framework based on qMRI maps that can describe the contribution of individual lipids to the membrane environment.

The human brain is comprised mainly (70-80%) of water, with proteins (8-11%) and lipids (5-15%) where the distribution of these molecules varies both between brain regions and across the lifespan, as well as in various pathological states^10–14^. There are five major groups of lipids: phospholipids (main component), cholesterol, and three types of glycolipids, cerebrosides, gangliosides, and sulfatides. Most of the lipids in the human brain are found in cell membranes where they are involved in many important biological circuits, and cellular functions^15^. Lipids are known to strongly affect the contrast of brain’s qMRI maps, a claim recently supported by experiments using lipid clearing techniques^16^.

Early NMR studies indicated that myelin, which is mainly composed of lipids, has a strong effect on image contrast and specifically to the T1 and MT contrasts. This was initially attributed to the interaction between water protons and cholesterol in myelin membranes^17^, although later studies showed that other lipids and macromolecules, such as proteins also influence the water proton magnetic relaxation^18–20^. Additional studies focused on the sensitivity of qMRI parameters to the lipid bilayer structure, indicated the influence of different surface groups and pH^21–23^. However there remains a great need for a systematic quantification of the effect of lipids on the qMRI parameters, in particular, while also monitoring the WF of the measurements.

This study describes a phantom system designed to assess the contribution of various membrane lipids to qMRI parameters. This system is based on an earlier version^24^ which we have now validated for reliability and generalized to increase the number of lipids used, the acquisition of qMRI maps, and the estimation of the water fraction.

The prepared liposomes are composed of naturally abundant brain phospholipids^25^ and our results demonstrate that the lipids membrane qMRI parameters, PD, T1, T2, and MT can be estimated with high reliability and accuracy. We propose that this system can serve as a tool to examine the contribution of lipid membrane composition to qMRI parameters.

## Methods

### Phantom samples

#### Sample Preparation

Liposomes designed to model cellular membranes were prepared from phospholipids by the hydration-dehydration dry film technique^26^.

The following highly purified and lyophilized lipids were used in this protocol (one or more in each experiment): L-α-Phosphatidylcholine(PC), L- α -Phosphatidylinositol(PI) and Cholesterol (Sigma-Aldrich), DS-Phosphatidylethanolamine 18:0/18:0(PE), DO-Phosphatidylserine 18:1/18:1(PS) and E Sphingomyelin(Spg) (LIPOID).

The lipids were dissolved in ethanol, methanol and/or chloroform over a hot plate and vortexed. Next, the solvent was removed to create a dry film by vacuum-rotational evaporation. Dulbecco’s Phosphate Buffered Saline (PBS), without calcium and magnesium (Biological Industries) was added to the dry film in order to obtain a final sample of 25-30% of liposomes, and 70–75% water by volume, at a pH similar to physiological conditions (pH ~ 7.5). The sample was then stirred on a hot plate at 65° for 2.5 hours to allow the lipids to achieve their final conformation as liposomes. Samples were further diluted with PBS to obtain preparations with liposome to water ratios ranging from 5-30%, for a final sample volume of 1.5 ml in a 4 ml squared polystyrene cuvette, 12.5 × 12.5 × 45 mm (light path: 10 mm ± 0.05 mm) with a lid. Cuvettes were glued to a polystyrene box, which was then filled with ~1% SeaKem Agarose (Ornat Biochemical) and ~0.0005M Gd (Gadotetrate Melumine, (Dotarem, Guerbet)) dissolved in double distilled water (ddw). The purpose of the agar with Gd (Agar-Gd) was to stabilize the cuvettes, and to create a smooth area in the space surrounding the cuvettes that minimalized air-cuvette interfaces (Fig. 1). The properties of the box were important for the bias correction stage of the analysis (see T1 maps & bias correction).

**Figure 1:**
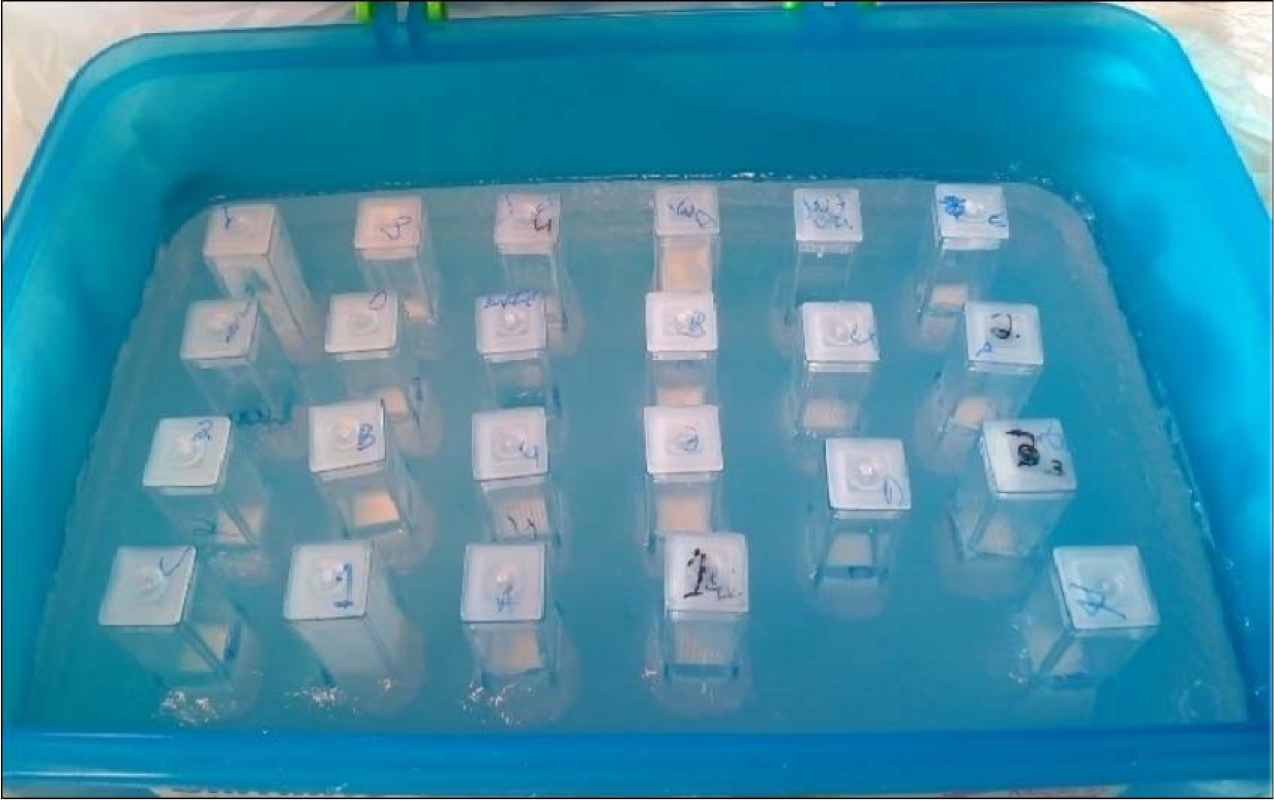
The Phantom box. 23 cuvettes filled with liposomes and water surrounded by homogeneous Agar-Gd gel inside a plastic container (22cm × 13 cm × 9 cm).

As controls, we also measured water based samples without lipids including PBS, ddw and a range of mixtures of ddw and deuterium (D_2_O).

#### Liposome Volume Estimation

We estimated the volume of 1 gram of a given phospholipid in water based on the molar mass and the combined volume of the head group and the tail (methylene groups) of a single lipid molecule. The volume of each lipid varies due to differences in the polar head groups, and the tail composition^27–29^. Here is a summary of the volume estimations used to calculate the lipid volume of each sample. For mixtures with cholesterol, we assumed the cholesterol volume to be negligible according to the free space theory^30^. The Estimated volumes of one gram PC/PC:Chol, PS, Spg and PI are 0.976 ml, 0.887 ml, 1.000 ml and 1.391 ml, respectively.

#### TEM and DLS measurements

Transmission Electron Microscopy (TEM) was used in order to verify and visualize the formation of liposomes. Samples of 10% liposome by volume were diluted by a factor of 10^2^, followed by centrifugation. The supernatant was harvested and diluted by a factor of 10. Final samples were negatively stained with uranyl acetate^31^.

In a separate experiment, the size distribution of the prepared liposomes was analysed by Dynamic Light Scattering (DLS,Malvern Zetasizer)^32^ on 10% samples diluted by a factor of 10^3^.

### Image Acquisition

The phantom system scan protocol was designed to resemble a human brain scan. Phantom samples were scanned in a 3 Tesla Skyra (Siemens) MRI scanner with a 32 channel receive head-coil. The phantom was kept in the magnet room for 20 min before the scan started in order to achieve thermal equilibrium. The magnet room temperature was kept constant at around 18°C.

#### Spin Echo Inversion Recovery (SEIR)

Single slice images were acquired using a *SEIR* sequence with adiabatic inversion pulse and inversion times of TI = 2000, 1200, 800, 400, 50, TR = 2540 ms, TE = 73ms, FoV of 222 mm, voxel size of 1.2 mm × 1.2 mm × 2 mm.

#### Spoiled gradient echo

Images were acquired using a FLASH sequence with 4 flip angles α = 4, 8, 16, 30, TE = 3.91 ms, TR = 18 ms, FoV of 205 mm, voxel size of 1.1 mm × 1.1 mm × 0.9 mm. The same sequence was repeated with a higher resolution of 0.6 mm × 0.6 mm × 0. 5 mm.

#### Spin echo

Images were acquired with multi spin echo sequence with 15 echo times TE = 10.5, 21, 31.5, 42, 52.5, 63, 73.5, 84, 94.5, 105, 115.5, 126, 136.5, 147, 157.5, TR = 4940 ms, FoV of 207 mm and voxel resolution of 1.1 mm × 1.1 mm × 1.1 mm.

#### Magnetization Transfer

Images were acquired with the FLASH Siemens WIP 805 sequence, TR = 72 ms, TE = 1.93, 4.46, 6.99, 9.52, 12.05, 14.58, flip angle = 6 deg, FoV of 205 mm, voxel size of 1.1 mm × 1.1 mm × 0.9 mm. An initial scan without an MT pulse was followed by 14 scans with RF offsets = [100, 200, 300, 400, 500, 600, 700, 800, 900, 1000, 2000, 4000, 50000, 10000 Hz], with MT flip angle of 220°. Repetitions with the same off-resonance pulses with MT flip angles of 540° and 760° were also performed for some of the experiments.

### Image processing

We used MATLAB to process the data acquired from the magnet. Manual segmentation of the areas of interest (ROI, agar, and sample cuvettes) was done using ITKsnap software^33^. Segmentation was designed to minimize both sample-plastic and sample-air interfaces. The median and the median absolute deviation were calculated for the ROI in each of the qMRI maps generated (below).

Code will be available at [Link to github will be added here].

### Quantitative Maps

#### T1 maps & bias corrections

The variable flip angle^34^ (VFA) approach (Eq. 1) for estimating T1 was selected as this is a commonly used technique for human brain quantitative T1 mapping.

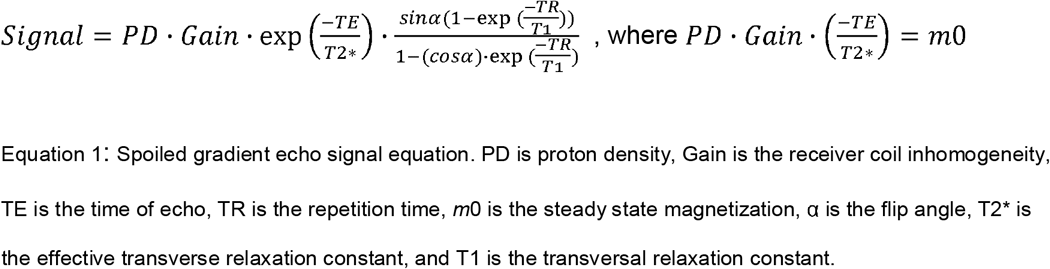

The VFA T1 mapping contains biases resulting from inhomogeneity in the excitation RF pulse. This inhomogeneity leads to spatial uncertainty regarding the flip angle used, i.e. the true flip angle. This error can be described as the percent difference between the nominal and the true flip angle value^35^ (Eq. 2)

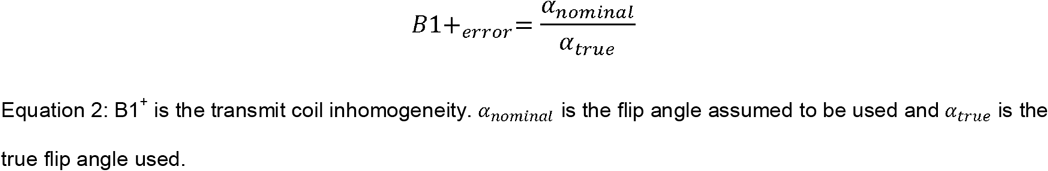

In order to account for the excite bias in the phantoms spoiled gradient echo T1 map, we modified previous methods^24,36^ to suit the phantom system, and combined them with a single slice Spin Echo Inversion Recovery (SEIR), which is considered to be a gold standard sequence for estimating T1 (GS-T1)^37^. This analysis is based on an adaptation of mrQ software^38^ and Vista Lab software^39^. The modification utilizes the fact that the Agar-Gd filling the box around the samples is homogeneous and can therefore be assumed to have a constant T1 value. We estimated the Agar-Gd T1 value using the algorithm reported by Barral et al.^37^and used this GS-T1 value to correct for the excite bias in the spoiled gradient echo space.

The analysis pipeline is described below:

1. We manually segmented the Agar-Gd in the SEIR T1 map and averaged its T1 value across the measured slice. This GS-T1 value is assumed to be accurate and free of biases.
2. We manually segmented the Agar-Gd in the 3D spoiled gradient echo volume.
3. We fixed the bias free GS-T1 value along the Agar-Gd ROI in the 3D spoiled gradient echo volume, and used Eq. 1 and Eq. 2 to calculate the flip angle percentage error numerically. This produced an estimate of the B1^+^ error values in each Agar-Gd location in the spoiled *gradient echo* volume.
4. Assuming B1^+^ is smooth in space, we interpolated and extrapolated the B1^+^ values to the whole volume and generated a full B1^+^ map using a low order polynomial^13^.
5. Using the B1^+^ bias map, we used Eq. 1 to calculate excite-bias free T1 and M_0_ values for the whole volume, including the cuvette samples.

#### Water Fraction maps

In order to evaluate the water fraction (WF) in the phantom system, we first used Eq. 1 to estimate the proton density (PD) from M_0_. Assuming that the M_0_ is proportional to PD and the receive coil gain inhomogeneity (Eq. 2), we can remove the spatial inhomogeneity and normalize the PD with a pure water sample.

To remove the receive coil gain inhomogeneity (g) from M_0_, and to calculate WF, we modified an existing method^24^ to suit the phantom system according to the following steps:

1. Assuming that the Agar-Gd’s PD values are homogeneous across space, any smooth variation detected in the Agar-Gd ROI in the phantom must be due to receiver inhomogeneity. The estimated bias was calculated locally in the Agar-Gd ROI by using low order polynomials. These local bias estimations were combined to obtain the inhomogeneity map over the whole volume.
2. The PD was obtained from the M_0_ and the calculated receiver inhomogeneity map using Eq. 1 (PD = M_0_/Gain).
3. In each phantom scanned, we included several pure water/buffer sample cuvettes. These samples were used to calibrate the PD map, such that the average PD values of the water cuvettes equal one. Applying this calibration to the PD map, allowed us to estimate a WF map where the samples are normalized to a WF between zero and one.

To calculate the non-water lipid fraction (equivalent to the term MTV in the human brain^24^), we subtracted the WF value of each sample from the WF value of pure water (i.e. MTV=1-WF).

To approximate T2* residuals on the PD estimations (see Eq. 1), we used spoiled gradient echo scans with multiple TEs.

With fixed values for all the signal equation parameters (i.e. T1, B1+ and Gain), the apparent PD (PD*) could be estimated as a function of the Signal at each TE according to Eq 3.

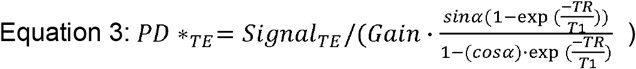

The measured signal could be extrapolated to TE = 0 (no contribution of T2*) using a second order polynomial model^40^. The value for WF value was obtained by normalizing the extrapolated PD value of the lipid samples to the extrapolated PD value of the water samples.

#### T2 maps

The T2 relaxation constant was estimated from a multi spin echo sequence by the EMC algorithm^41^. This algorithm assumes that the actual signal received is not a perfect decaying exponent and obtains the result by simulating the Bloch equation and utilizing a lookup table of the parameters used in the sequence.

#### MT Maps

A spoiled gradient echo sequence with multiple off-resonance RF pulses was used in order to estimate the ratio of MT_on_/MT_off_; (termed MT_norm_ for simplicity)^22^. Having compared the data for three MT off-resonance flip angles, we chose to work with the MT flip angle of 220° and the 700 Hz off-resonance pulse, where the direct effect of water appears to be minimized. This can be seen in Fig. 2, where the Z spectrum of water reaches a value of approximately one at 700 Hz.

**Figure 2:**
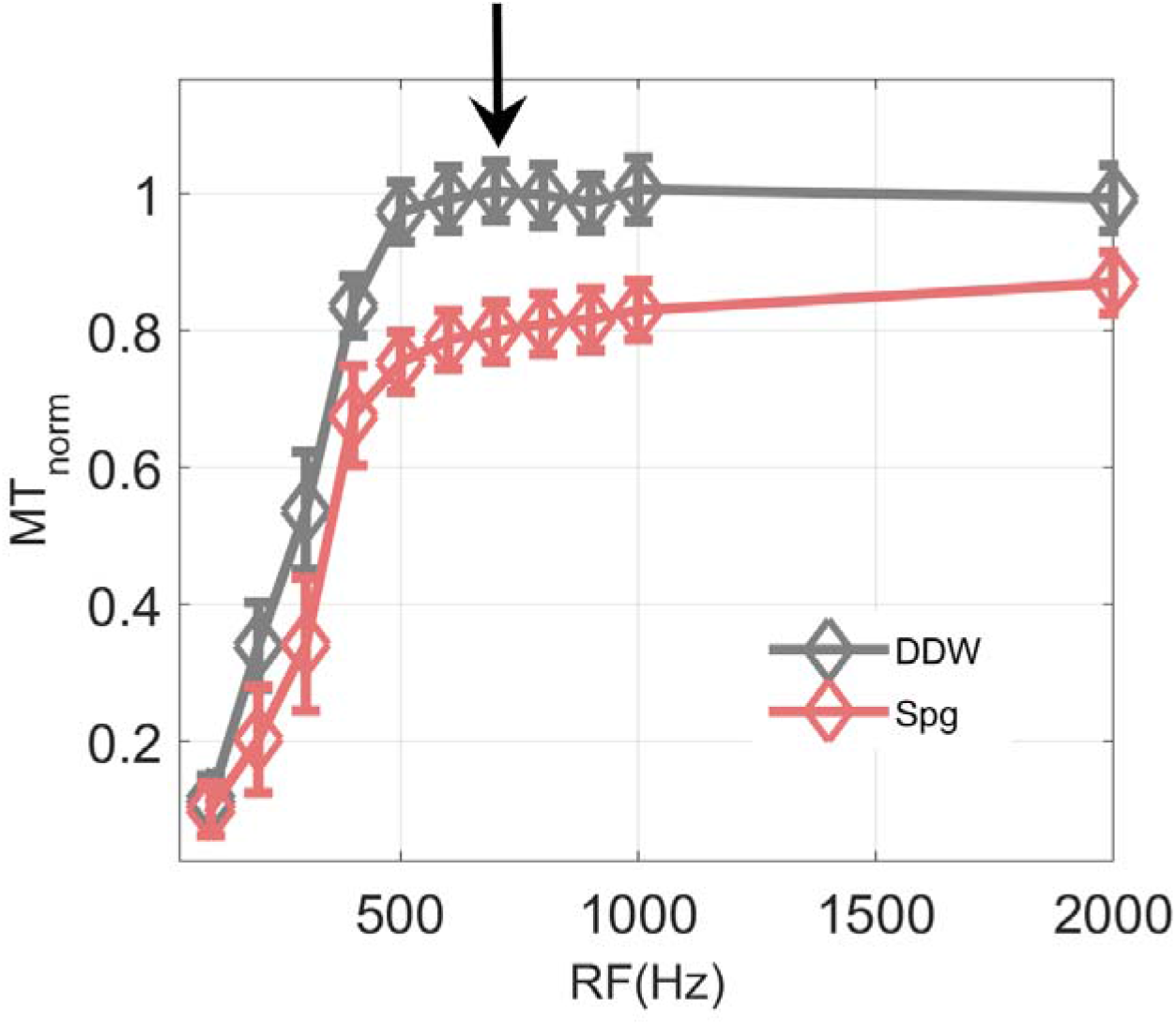
Examples of MT_norm_ Z-spectra for water and a liposome sample of sphingomyelin (Spg). Arrow indicates an RF pulse of 700Hz. The error bars represent the STD for each RF pulse in the relevant sample.

## Results

This study evaluated a phantom system involving liposomes as a tool for studying the effect of biologically relevant membrane lipids on qMRI measurements.

Figure 3a,b shows TEM images of the liposomes, with the lipids organized in spherical membrane structures as expected, indicating the success of the manufacturing process. A typical size distribution of the liposomes with an average diameter of 1-10 μm is shown in Figure 3c.

**Figure 3:**
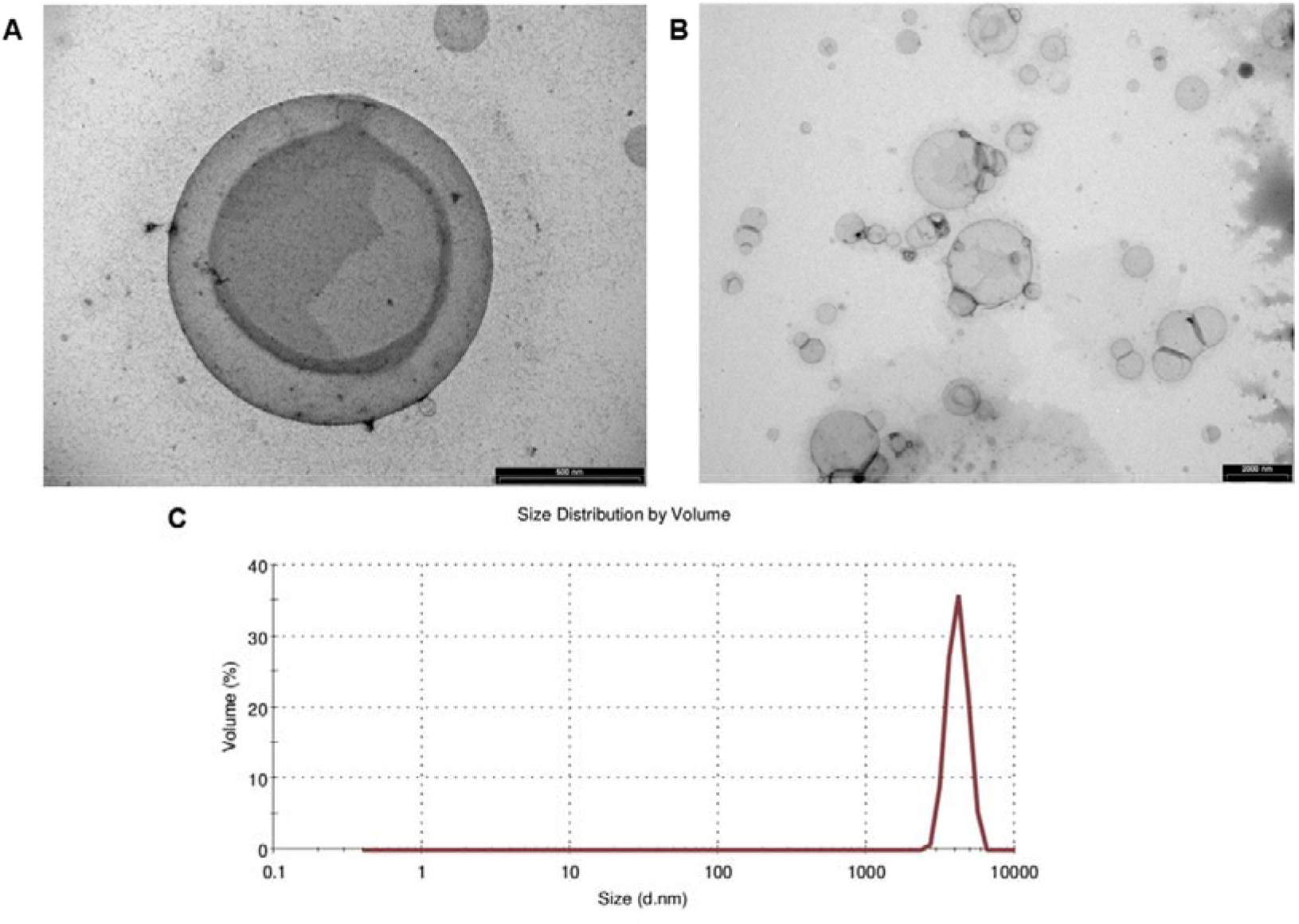
(A:B) TEM images of liposomes. Left panel center shows a single liposome about 1μm in diameter. This liposome is most probably comprised of two layers of phospholipids shown by two distinct darker black borders. Right panel: a wide distribution of liposome sizes. Overlaps between liposomes are mostly due to the imaging technique (C) Example of liposome size distribution from DLS measurements of different samples.

### Establishing an experimental system for qMRI of the phantom

#### Homogeneity and stability of the Agar-Gd

The liposomes samples are contained in plastic cuvettes that are embedded in agar with Gd ions (Agar-Gd). The Agar-Gd ROIs are used for the estimation of both receive and transmit biases by assuming the ROIs creates an envelope of homogeneous area. Figure 4 shows an example of a slice through the phantom array (Fig. 1) presenting maps of the qMRI parameters PD, T1, T2 and MT_norm_.

**Figure 4:**
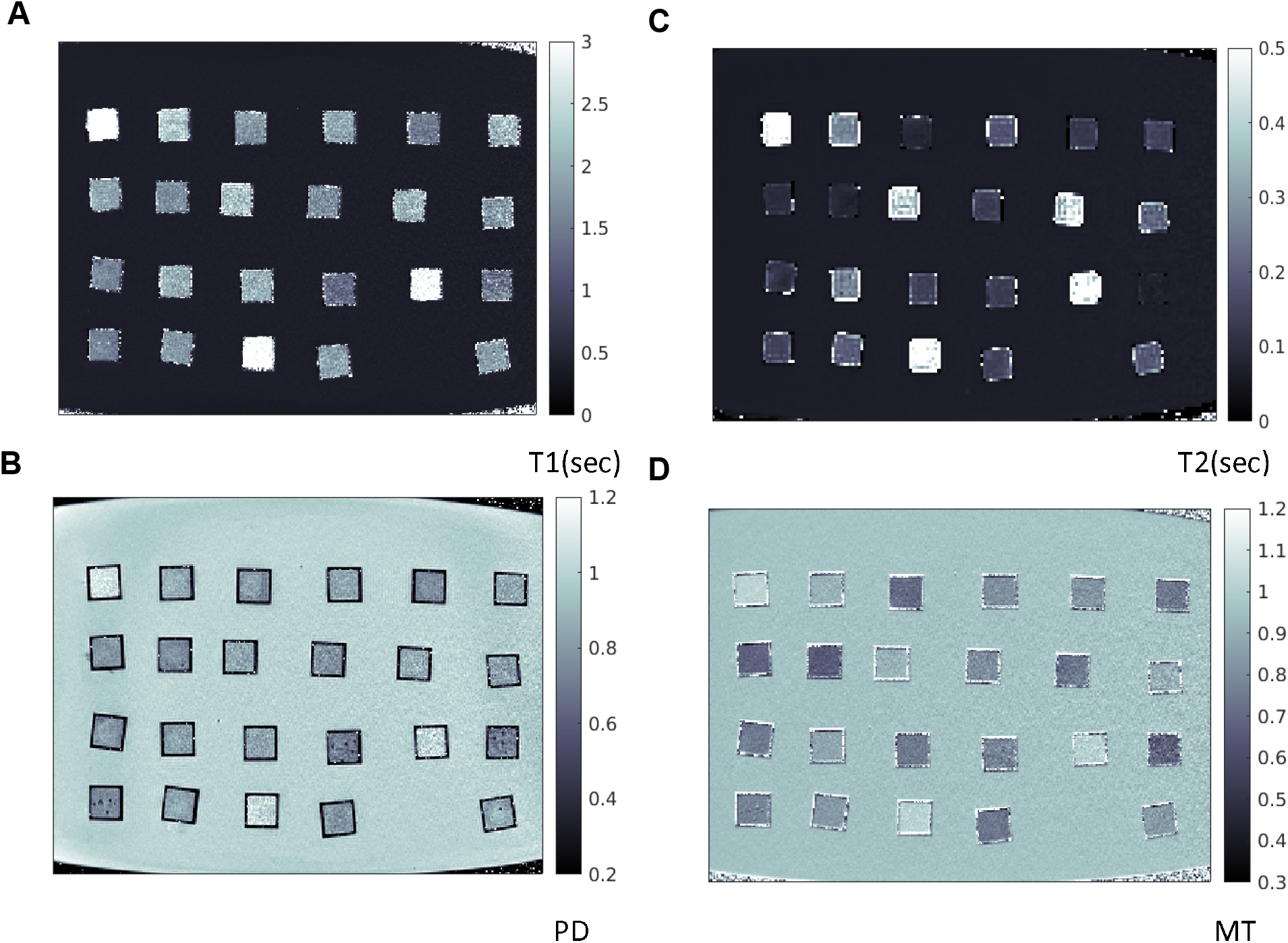
Quantitative MRI maps. (A-D) A slice through the phantom array shown in Figure 1 indicating the T1, PD, T2, and MT_norm_ maps respectively. The cuvette borders can be seen in black (Figure 4B). The Agar-Gd gel area around the cuvettes is a relatively constant area. Each cuvette has different values reflecting the liposome content and water fraction.

GS-T1 analysis indicated that the Agar-Gd ROI was indeed homogeneous and gave stable results in space (up to 3.5% change; N = 31) and time (N = 31 different days) with T1 values (421 ± 63 ms).

Importantly, while our corrections are calculated only on the Agar-Gd ROI, they can then be extended to correct for the total volume of the system, which includes the lipid sample ROIs. Figure 5a shows a typical distribution of T1 values for Agar-Gd before and after the B1^+^ correction and Figure 5b presents a typical B1^+^ bias map. In addition, a typical receiver gain bias is shown in Figure 6a, while Figure 6b displays a corrected PD map. These corrections are designed to remove the biases from the T1 and PD estimations. Summarizing multiple experiments, the average STD of the Agar-Gd ROI in space was 2.7% for VFA-T1 and 1.7% for WF. Notably, the EMC T2 mapping algorithm, which includes a build-in bias correction, generated a stable T2 in space (median value of 0.07 sec, N = 30), with a typical STD of 2% of the signal. Similarly, the MT_norm_ parameter, which corrects for the transmit bias with the MT_on_ to MT_off_ division, had a small STD of 1.75% in space (median value of 0.95, N = 30). Repeated measurements of the relaxation of water in multiple experiments, showed high consistency (T1 of 3837 ± 892 ms, T2 of 0.66 ± 0.05, MT of 0.99 ± 0.02). We therefore suspect that some of the variability can be attributed to small temperature differences (1-3°C)^45^.

**Figure 5:**
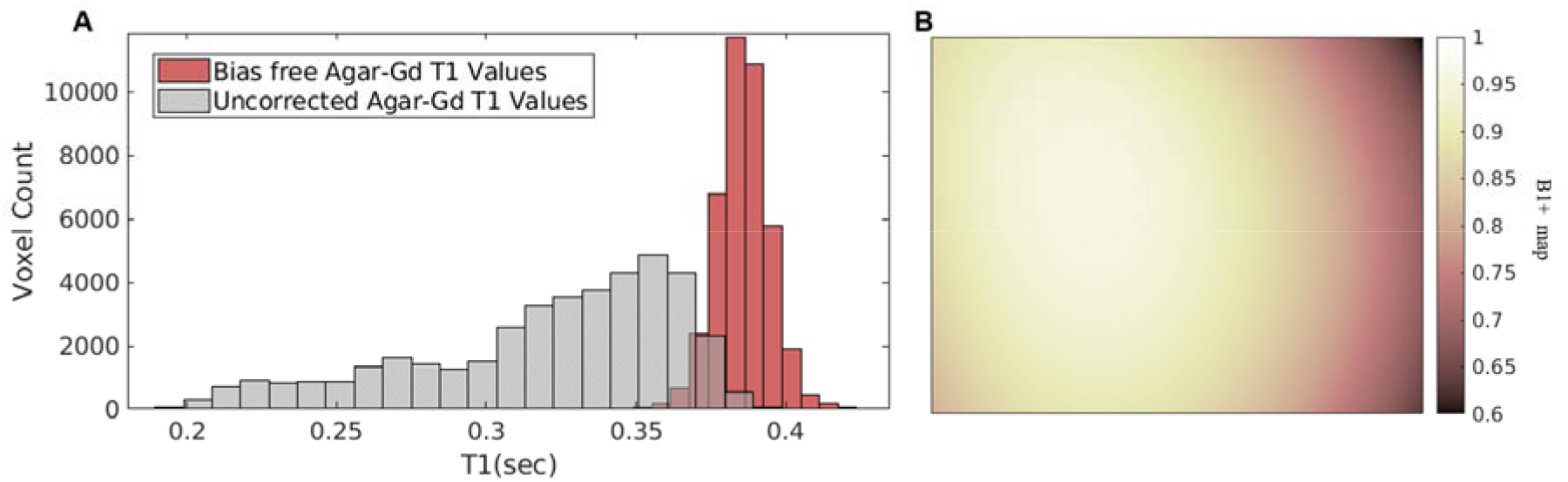
(A) Distribution of T1 values estimated by the VFA technique before and after the B1 + correction. (B) B1+ map. A B1^+^ inhomogeneity is represented by a typical peak in the center of the imaging volume (brighter shades in the center and darker shades in the periphery).

**Figure 6:**
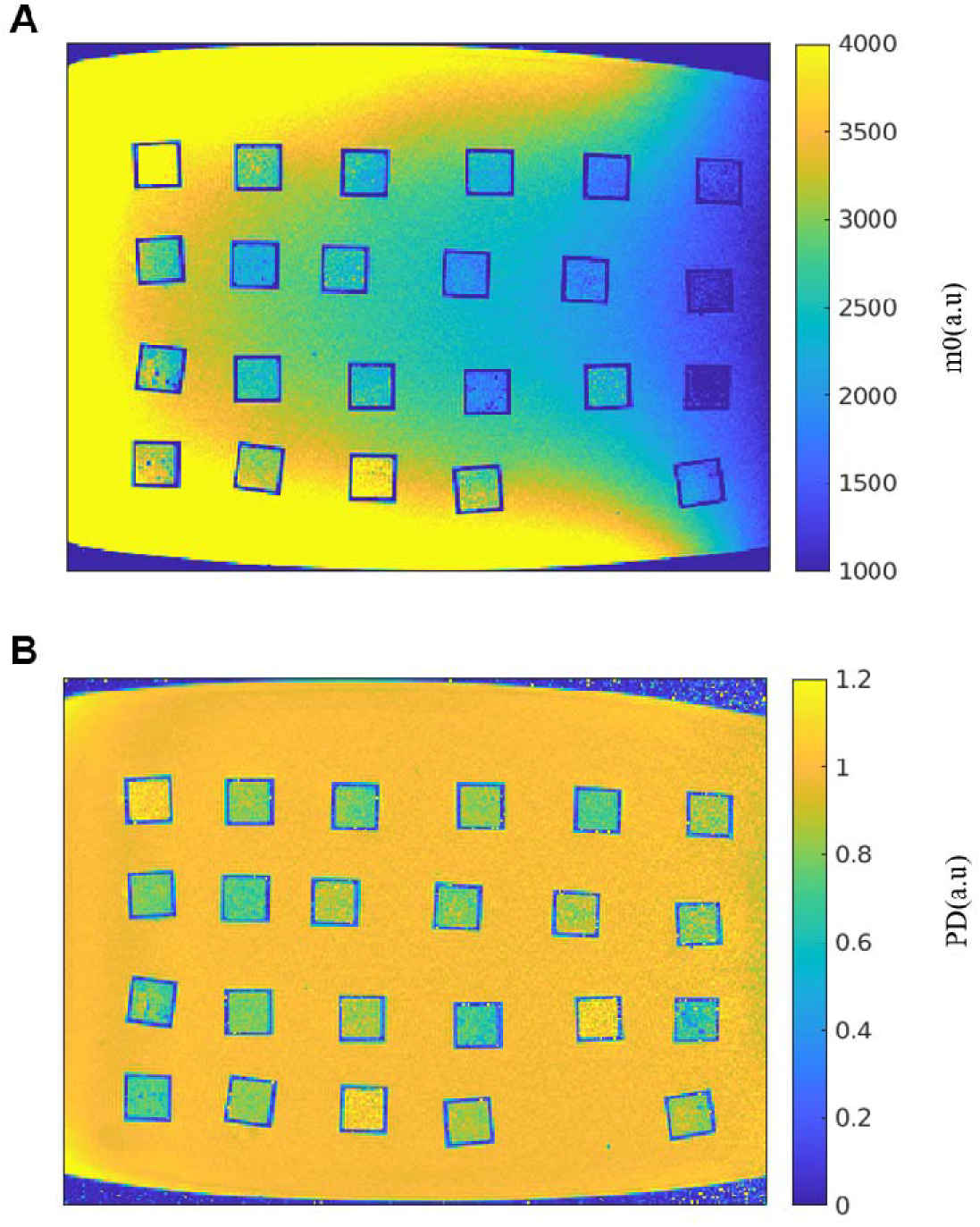
(A) M_0_ map. The receive coil gain inhomogeneity can be seen across the image (yellow to blue shades). (B) PD map after the removal of the gain of the receiver coils.

#### Homogeneity, stability and consistency of the liposomes

To test the reliability and stability of the qMRI liposome phantom measurements, we compared multiple scans of the same liposome samples taken on different days. We found remarkable consistency in T1, T2, MT_norm_ and WF estimations across scans (Figure 7), with a high coefficient of determination (R^2^ = 0.90, 0.93, 0.85, and 0.93 respectively, N = 78). Next, we tested the distribution of values within a sample ROI. There was no slice profile within the cuvette along the Z direction (the XY plane is narrow in comparison to the long Z axis of the cuvette) (Figure 8). We used a one-way Anova test to rule out any outlier slices in space. We found no significant differences between slices of cuvettes from randomly selected sets of liposomes in different experiments (mean P-value of 0.43 ± 0.3 with mean STD of 0.3, N = 32).

**Figure 7:**
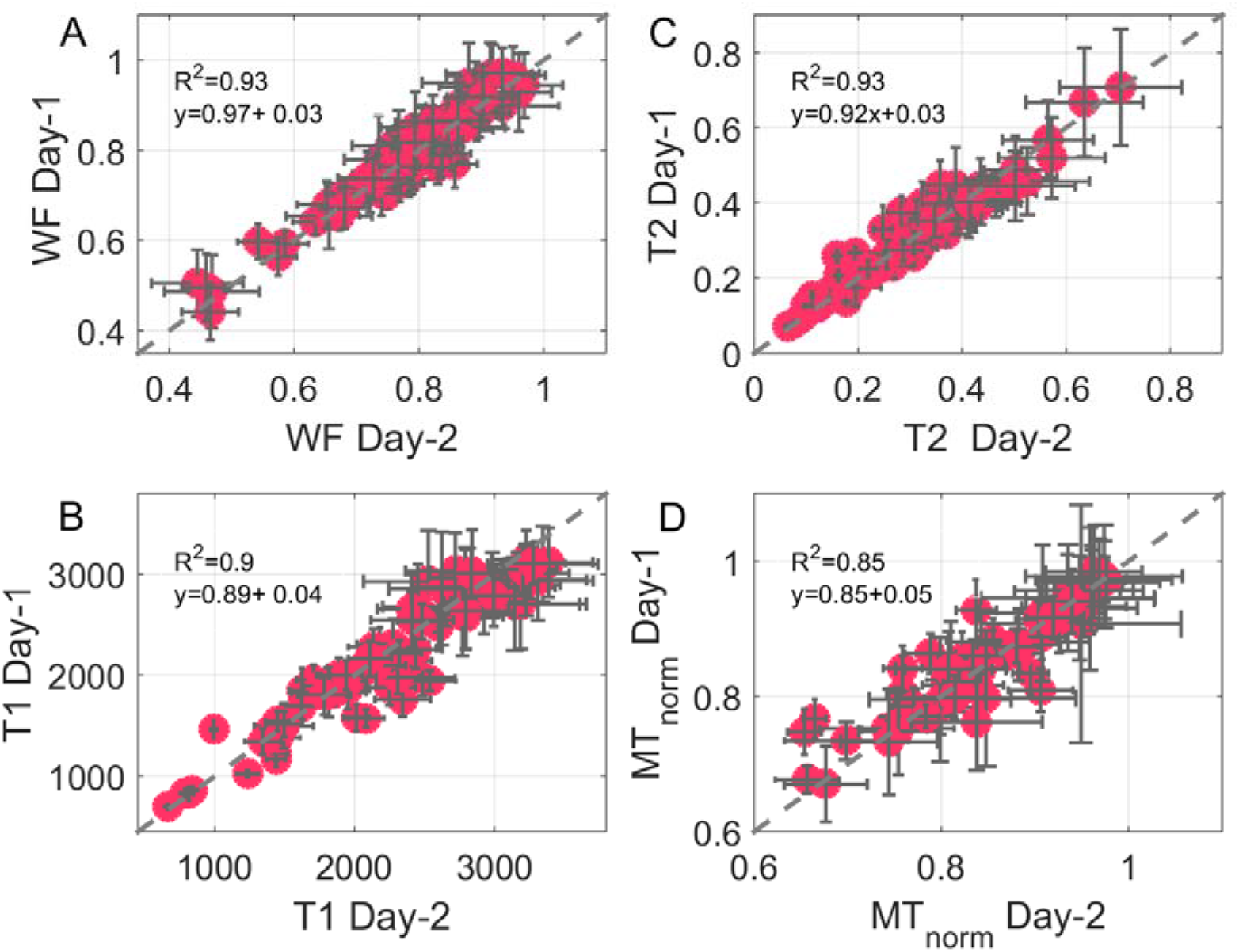
Reliability of the qMRI parameters. (A:D) Relaxation constants were fitted for two scans of the same lipid sample on different days. Panels A:D represent: WF, T1, T2 and MT_norm_, respectively. MT_norm_ is the MT offset at 700 Hz, flip angle of 220 degrees (where the direct effect is minimalized) normalized by the same scan without MT. Values estimated on different days were very reproducible. Error bars corresponds to median absolute deviation.

**Figure 8:**
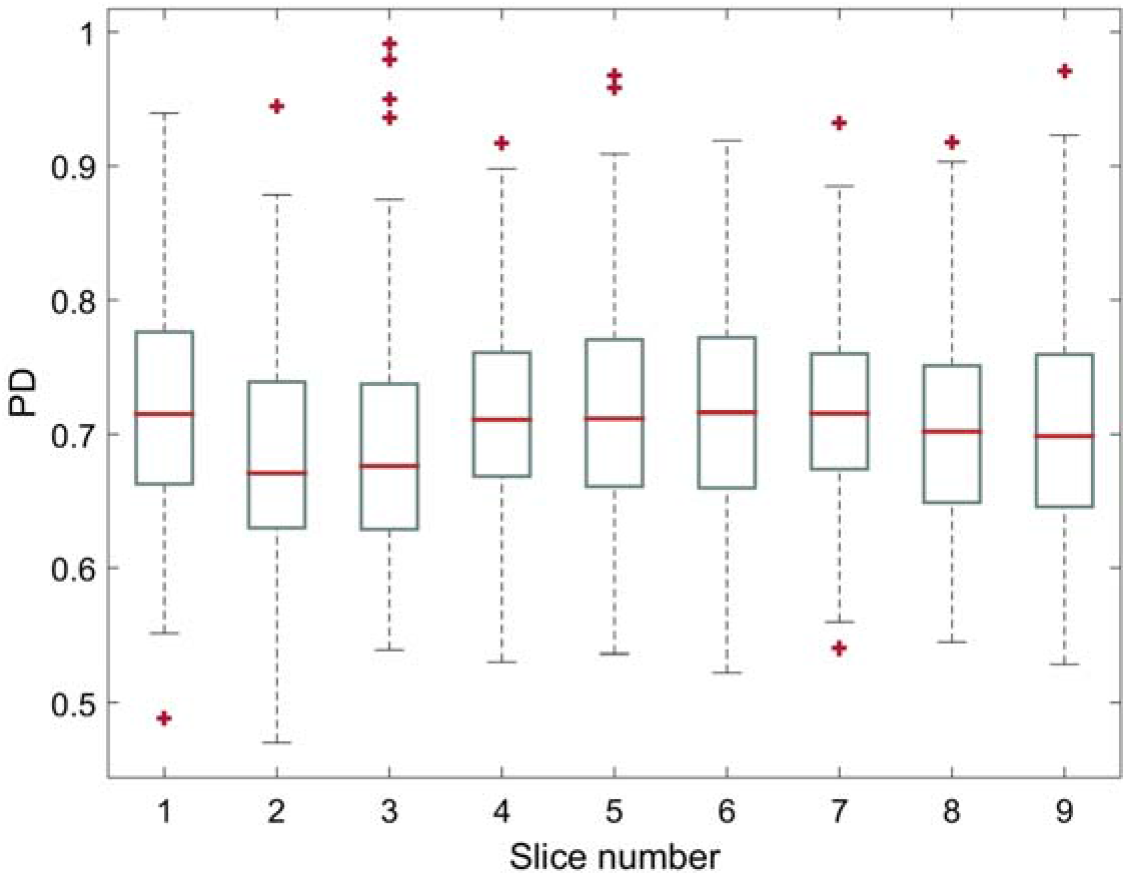
Profile of PD values across slices of a single liposome-containing cuvette.

### Reliability of the WF estimations

In order to validate the accuracy of our WF estimation, we created a line of samples with varying ratios of H_2_O to D_2_O. Since deuterium is an MR-invisible isotope, H_2_O/D_2_O mixtures can be used to calibrate water fraction estimations^42^. Our results indicated an excellent agreement between the WF and the true volume fraction (R^2^ = 0.99, values are found on the identity line, Figure 9a).

**Figure 9:**
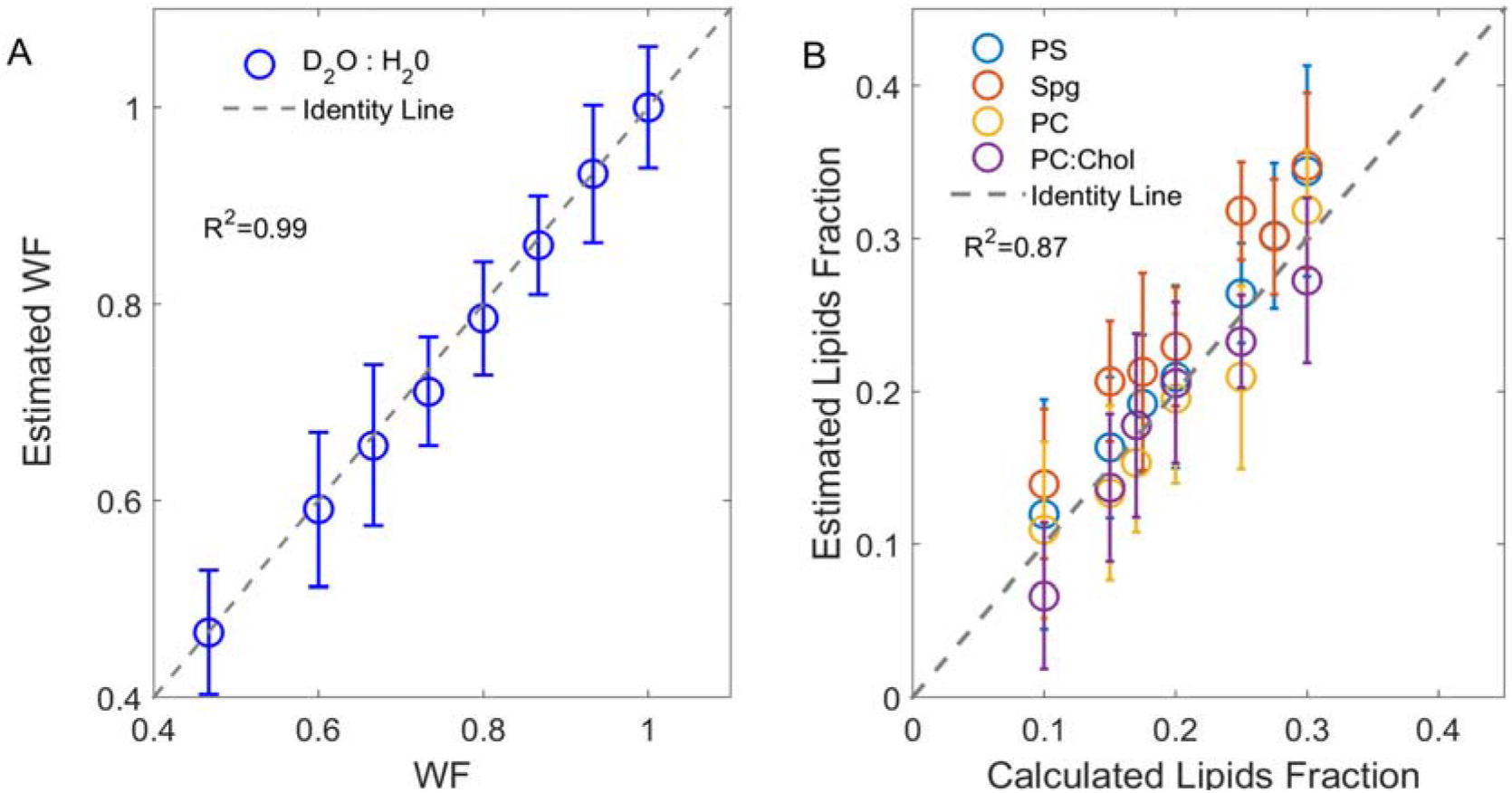
Accurate water and lipids fractions estimation. The MRI based WF estimation (y-axis) agrees well with the true WF (x-axis). Each point represents the mean and standard deviation of all the voxels within a different sample. (A) Each sample is composed of different ratios of H_2_O (H-MRI visible) and D_2_O (H-MRI invisible). (B) Lipid fraction (non-WF) estimations of sample with liposomes having different lipids volume fractions. The true lipid fraction (x-axis) was determined using the theoretical volume of the lipids molecules taken from the literature, and agrees well with the qMRI-based estimated non-WF(1-WF) (y-axis). The theoretical WF estimation does not account for water loss during the manufacturing process.

An analytical estimate of phospholipid volume provided an additional opportunity to verify the estimations of WF. The calculated lipid volume (see Methods: Liposome Volume Estimation), showed a good agreement (R^2^ =0.87, Figure 9b) to the qMRI-based estimated lipids fraction (the non-water fraction), which was defined as 1-WF (equivalent to the term MTV in the human brain^24^).

As a final verification, we tested our assumption that the short TE used (3.9 ms) did not introduce any T2* contributions that confound the estimation of WF in the lipid samples in our system. First, we compared the WF estimations with TE of 3.9 ms, to the WF estimation with the shortest measured TE (1.9 msec) and found only a negligible difference (Figure 9a, R^2^=0.86, values dispersed around the identity line with a slope of 0.95). Additionally, to remove any residual T2* contribution to the WF value, we used multi-TE data and fit a second order polynomial in order to extrapolate^40^ the WF value at TE = 0. Here again we found a negligible effect of TE dependency on the WF estimations (R^2^ = 0.58, values dispersed around the identity line, with a slope of 0.81, Figure 9b). Altogether these results demonstrate a negligible T2* effect given the liposome sample content and the short TE used in our experiments.

## Discussion

Quantifying the contribution of lipids and macromolecules to the qMRI signal is key for the understanding of in vivo human imaging. This study describes the characterization of a biologically relevant liposome phantom system. The non-water volume fractions in our samples were 5%-40%, which are similar to the brain’s grey and white matter values. The use of this liposomes phantom system enables us to control the lipid composition and to reliably measure the contribution of various lipids to qMRI parameters.

An important feature of the system is the homogenous Agar-Gd surrounding the cuvettes, which effectively corrected the qMRI measurements of the liposome samples for the different spatial inhomogeneities in VFA scans. After correction, the distributions of values within the samples was not influenced by slice profiles, indicating that our liposome samples are homogeneous, with minimal amounts of unwanted precipitates or air bubbles. Repeated scans and experiments revealed that the STD in our system are at least one order of magnitude smaller than the values of the estimated parameters. In some cases, we identified minor variations that may be due to temperature fluctuation, changes in the structure of the liposomes with time, and imperfections in post-processing, such as masking of non-identical voxels. Interestingly, samples with higher WF had higher variability (Figures 7) as described previously in the literature^36^.

This study employed the MT_norm_ convention, with 700 Hz off-resonance frequency selected as our MT_on_ signal. Since water samples at ~700 Hz with MT power flip angle of 220° gave a value of one in our system, we could estimate the MT component in our samples by minimizing the direct effect of water.

For T2 mapping, we used the EMC approach, and verified that this fitting technique generated reliable and bias-free T2 maps.

Developing this protocol will allow us to accurately measure the qMRI parameters in a lipid environment as verified by the stability of the qMRI parameter results from multiple repetitions and scans done on different days. Another important aspect of this phantom system is the ability it gives researchers to control and change the size distribution of the liposome samples by the use of sonication^32^. Our findings suggest that this phantom system can be generalized and used to compare lipids manufactured and scanned at different times (Figure 6). While the experiments presented here employed settings very similar to those used for human qMRI scans^13^, we suggest that a similar phantom could also be used to test a variety of qMRI models^43^ and novel qMRI techniques^44,45^.

The quantitative WF maps prepared in the study were validated as reflecting the true water and lipids volume fractions of the prepared samples (Figure 9). In the case of the liposomes, the T2* contributions to the WF estimation were negligible (Figure 10). The accuracy of these WF estimations allow us to fully rely on the qMRI measurements and mean that this estimation method could be useful for other biological samples with unknown WF.

**Figure 10:**
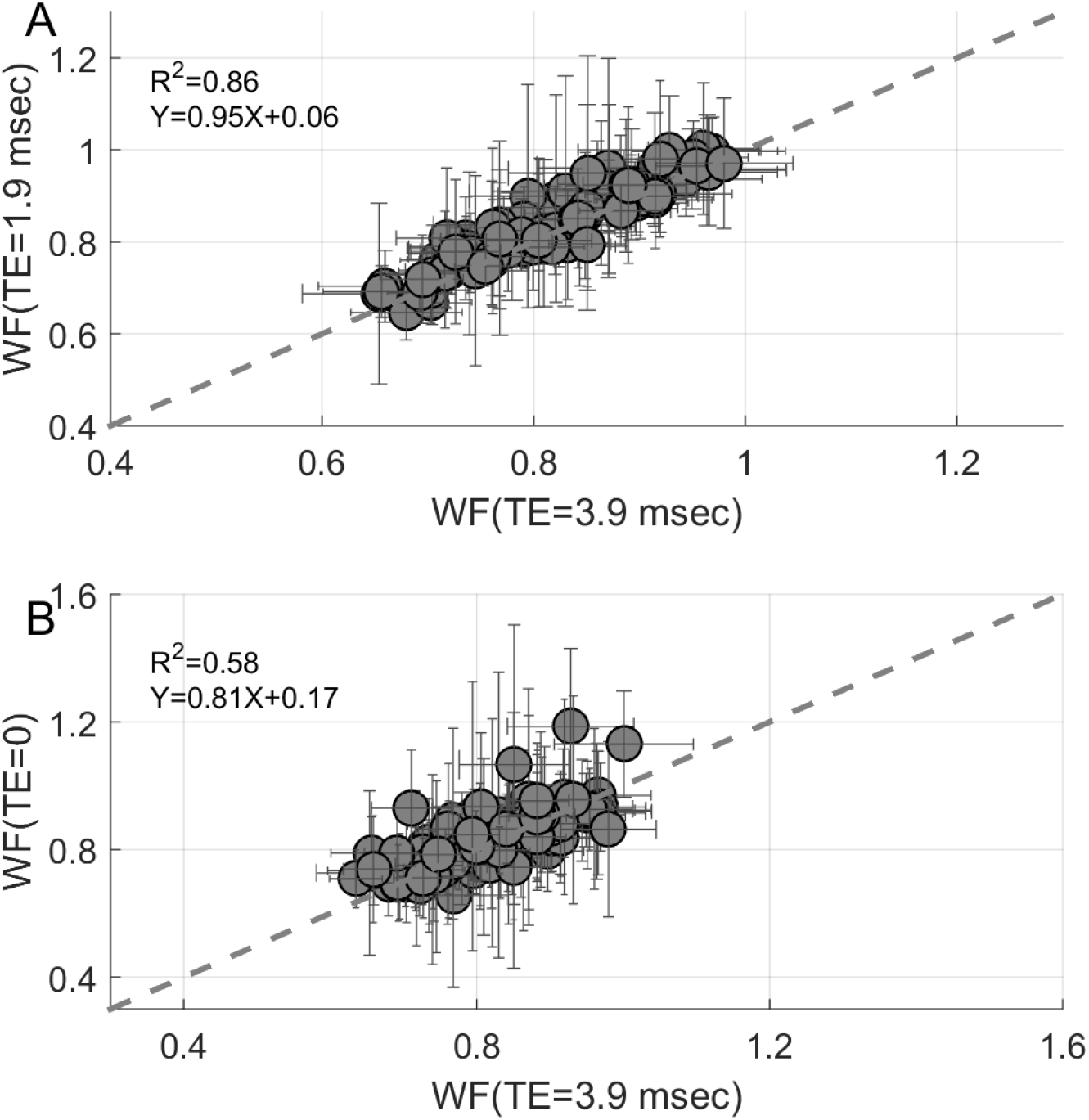
T2* effect on WF estimation. (A) Accounting for T2* contribution to WF estimations from a VFA-T1 map of liposomes. The possible effect of T2* on WF is calculated from a multi-echo scan. Each point represents the WF and median absolute deviation of the sample. Comparison between the estimation of WF by the VFA approach (TE = 3.9 ms) to WF values calculated using multiple TE values (TE = 1.9 ms). (B) Comparison between the estimation of WF by the VFA approach (TE = 3.9 ms) to multiple TE values and extrapolation to TE = 0. In both cases WF values are around the identity line with negligible dependency on the TE.

Establishing a stable but flexible MRI phantom system for lipids has great potential for the investigation of the contribution of lipid composition to qMRI parameters. In particular, this will allow us to extend early NMR findings, which suggested that relaxation depends on the magnetic field strength, temperature, and pH^22^.

Moreover, our liposome system provides a useful tool to model how water relaxation is influenced by different molecular environments. A key point of this system is the estimation of WF together with other qMRI parameters. Importantly, it was argued that this information may disentangle the WF contribution to the qMRI parameters from the physical-chemical contributions^1,24^.

In conclusion, we have successfully developed a biologically relevant liposome phantom system with controllable content that enables the measurement of the contribution of various lipids to multiple qMRI parameters in a reliable and accurate fashion.

## Acknowledgments

This work was supported by the ISF Grant (no. 0399306) awarded to A.A.M. We thank Batsheva Weisinger and Jonathan Bain for helpful comments on this manuscript. We thank Dr. Adi Pais for her help in designing the scan protocols. We thank Joohoon Ha for his help with the experiments.

